# Response reversal during top-down modulation in cortical circuits with multiple interneuron types

**DOI:** 10.1101/124669

**Authors:** Luis Carlos Garcia del Molino, Guangyu Robert Yang, Jorge F. Mejias, Xiao-Jing Wang

## Abstract

Pyramidal cells and interneurons expressing parvalbumin, somatostatin, or vasoactive intestinal peptide show cell type-specific connectivity patterns leading to a canonical microcircuit across cortex. Dissecting the dynamics of this microcircuit is essential to our understanding of the mammalian cortex. However, experiments recording from this circuit often report counterintuitive and seemingly contradictory findings. For example, the response of a V1 neural population to top-down behavioral modulation can reverse from positive to negative when the bottom-up thalamic input changes. We developed a theoretical framework to explain such response reversal, and we showed how this complex dynamics can emerge in circuits that possess two key features: the presence of multiple interneuron populations and a non-linear dependence between the input and output of the populations. Furthermore, we built a cortical circuit model and the comparison of our simulations with real data shows that our model reproduces the complex dynamics observed experimentally in mouse V1. Our explicit calculations allowed us to pinpoint the connections critical to response reversal, and to predict the existence of more types of complex dynamics that could be experimentally tested and the conditions to observe them.

## Introduction

Three major non-overlapping classes of interneurons expressing parvalbumin, somatostatin, or vasoactive intestinal peptide (henceforth denoted PV, SST and VIP respectively) make more than 80% of GABAergic cells of mouse cortex [28]. These neurons show cell type specific connectivity within themselves and with excitatory (E) neurons [9, 25] leading to a canonical microcircuit in cortex. There has been a lot of interest on the function of interneurons [6, 11, 13–15, 17, 31, 32], however we still do not fully understand the mechanisms that underlie the behavior of this microcircuit which are often complex and counterintuitive.

One particular example of complex behavior is the modulation of responses to visual stimuli during locomotion, when V1 activity significantly increases with respect to immobility [22] even in complete absence of visual input [10]. VIP interneurons are known to be involved in such modulation because artificially activating (damaging) them mimics (blocks) the effect of running on visual response [4]. Since VIP cells inhibit SST cells which in turn inhibit excitatory, PV and VIP cells, a natural explanation for this phenomenon is disinhibition [16, 31]: upon activation of VIP cells the SST population is inhibited and therefore neurons targeted by the SST population are disinhibited, raising the overall rate of the excitatory neurons. However recent experiments show that the network behavior might be more complex. In particular in the absence of visual stimulation, the activation of VIP cells results in an average decrease of SST population activity [3, 4] whereas in the presence of visual stimulation the response of SST cells is reversed and its rate increases during locomotion [3, 24] which appears to challenge the disinhibition hypothesis. This observations suggests that the nature of the interaction between VIP and SST could be stimulus dependent.

These experimental results raise two questions: First, the external activation of a population that directly inhibits a second population can trigger a positive response of the latter. What is the mechanism behind this apparently paradoxical behavior? Second, the same top-down modulation can trigger both a positive or a negative response of certain populations of the circuit depending on the sensory input. Under which conditions can we expect one response or the other?

In this study we model cortical activity and provide a comprehensive explanation to these two questions. We show that these counterintuitive phenomena rely on two basic features of cortical networks: (i) the presence of multiple populations of interneurons and (ii) nonlinear responses to input. Our framework is general and we use it to predict complex behaviors that have not yet been experimentally tested.

## Results

We simulate microcircuit activity using a four population firing rate model. The average rate of each population is given by a nonlinear function of its input that we refer to as the f-I curve [1]. The f-I curve is such that when the input is low (below threshold) cells are little responsive to changes in external input. Instead for high input (above threshold) small changes in the input can drive substantial changes in the response. This nonlinearity has been analyzed experimentally and theoretically [21, 26] and as we will show later, it is a key feature of the model.

Populations are connected according to the microcircuit scheme in (figure 1a) which contains the connections reported in both [9] and [25]. We also consider three sources of input: (i) top-down modulation that targets VIP cells (ii) local recurrent input and (iii) constant background input set so that the populations have some xed baseline activity (see methods for details).

**Figure 1:**
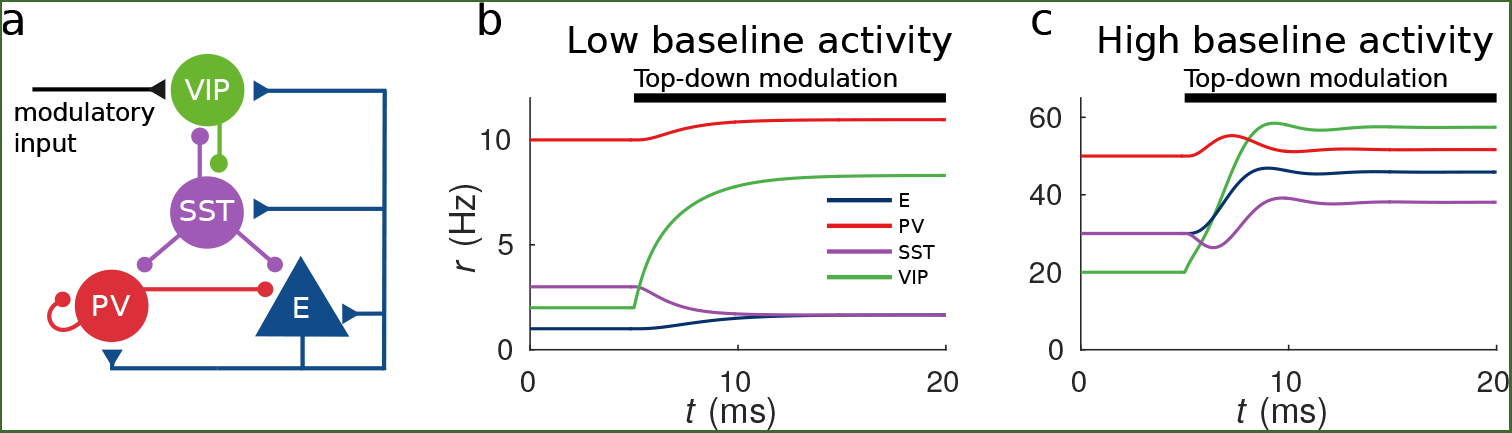
Response to top-down modulation depends on baseline activity. (a) Microcircuit connectivity and top-down modulatory input. (b, c) Transient dynamics upon the onset of the top-down modulatory current for low baseline activity (b) and high baseline activity (c). Under the low baseline activity condition SST is inhibited and E and PV are slightly disinhibited. The high baseline activity condition shows an example of response reversal in SST activity: it initially goes below the baseline rate but due to signi cant change in E activity and to the recurrent excitation it eventually reverses to a rate higher than baseline.

### Response to top-down modulation depends on baseline activity

To illustrate possible complex behaviors displayed by the network, we first focused on the circuit responses to top-down modulation. The simulation results from our model allow us to identify two qualitatively different scenarios. On the one hand, when the baseline activity of the network (i.e. activity before the onset of the top-down modulation) is low, the rate of the SST population decreases with respect to the baseline while the rates of the other populations (E, PV and VIP) increase (see figure 1b). On the other hand, when the baseline activity is high, the rate of all populations increases with top-down modulation (see figure 1c).

The surprising behavior exhibited by the SST population can be explained heuristically by analyzing the response of the different populations to external excitatory input targeting VIP cells. When the top-down modulation starts, the rate of the VIP population increases. This effect initially results in a reduction of SST activity and therefore a reduction of inhibition to VIP, PV and E cells. When baseline activity is low the E population is below threshold and this change in net input has a small effect in the output. In that situation all populations quickly reach a stationary state. However, when the baseline activity is high the E population is above threshold and a small change in input from SST cells has a big effect on the rate of the E population. If the recurrent excitation in the microcircuit is strong enough it can reverse the initial response of the SST population making it increase its activity to a higher rate than the baseline.

### Circuit behavior explained by response matrix

In order to formally characterize the steady state response of a population to external input we introduce the response matrix *M*. The intuition behind the response matrix is that if we change the input to population *j* (where *j* = *E*, *P*, *S*, *V* for excitatory, PV, SST and VIP populations respectively) by a small amount *δI_j_*, then the change in rate of the population *i* will be *δr_i_* = *δI_j_M_ij_*. If *M_ij_* _is_ positive (negative), an increase of the external excitation to *j* will result in an increase (decrease) of the rate of population *i* (see methods and table 3 for details). In contrast to the connectivity matrix, which takes into account only the direct path from population *j* to *i*, the response matrix contains information about all the possible ways in which population *j* can affect population *i*, namely through indirect connections *j*-*h*-*i*. Due to the complexity of these indirect pathways, for different values of the connectivity matrix (but preserving the excitatory/inhibitory structure) *M_ij_* _can be positive or_ negative irrespective of whether the connection from *j* to *i* is inhibitory or excitatory. Furthermore due to the nonlinearities in the f-I curve, the response depends on the baseline rate of each of the populations and, as shown before, it can reverse its sign.

As an example we analyze in detail the term

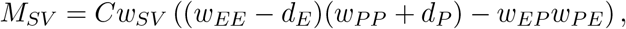

where *w_ij_* are the absolute values of the connection weights and therefore are positive by definition and for the system to be stable *C* has to be positive (see methods for details). The terms *d_i_* are proportional to the inverse of the first derivative of the f-I curves and are always positive. In particular *d_E_* becomes arbitrarily large when the input is very low and tends monotonically to a positive constant 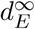 for high input. Therefore, if 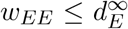 then *M_SV_* will always be negative. However, for 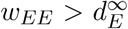 the behavior is much richer: if input is high then *d_E_* will be close to its minimum 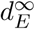 and *w_EE_* > *d_E_* allowing for *M_SV_* to be positive (provided that the product *w_EP_w_PE_* is small enough). Instead if the input is low, *d_E_* will become very large and *M_SV_* will be negative.

It is remarkable that this change in the interaction between VIP and SST populations depends on the activation level of E: modifying the state of one population has a impact in the interactions between other populations. The heuristic explanation is that if the recurrent excitation is strong enough and the E population is already strongly excited (above threshold), a small decrease in the inhibition from SST to the E population can boost its activity and therefore strongly drive the whole microcircuit. If instead, the E population is in a low activation state the change in inhibition will have a weak effect that will not be able to reverse the response of SST.

This observation provides an explanation to the reversal of the response of SST to VIP activation when the baseline activity is changed: as we show in figure 2a and 2c for low baseline activity, *M_SV_* _is_ negative and the presence of an external excitatory current targeting VIP cells will result in a negative response of SST cells and positive response of E, PV and VIP cells, conforming to the disinhibitory hypothesis. On the other hand, for high baseline activity (panels 2b and 2d), the response of the SST population to input to VIP cells becomes positive leading to the response reversal regime.

**Figure 2:**
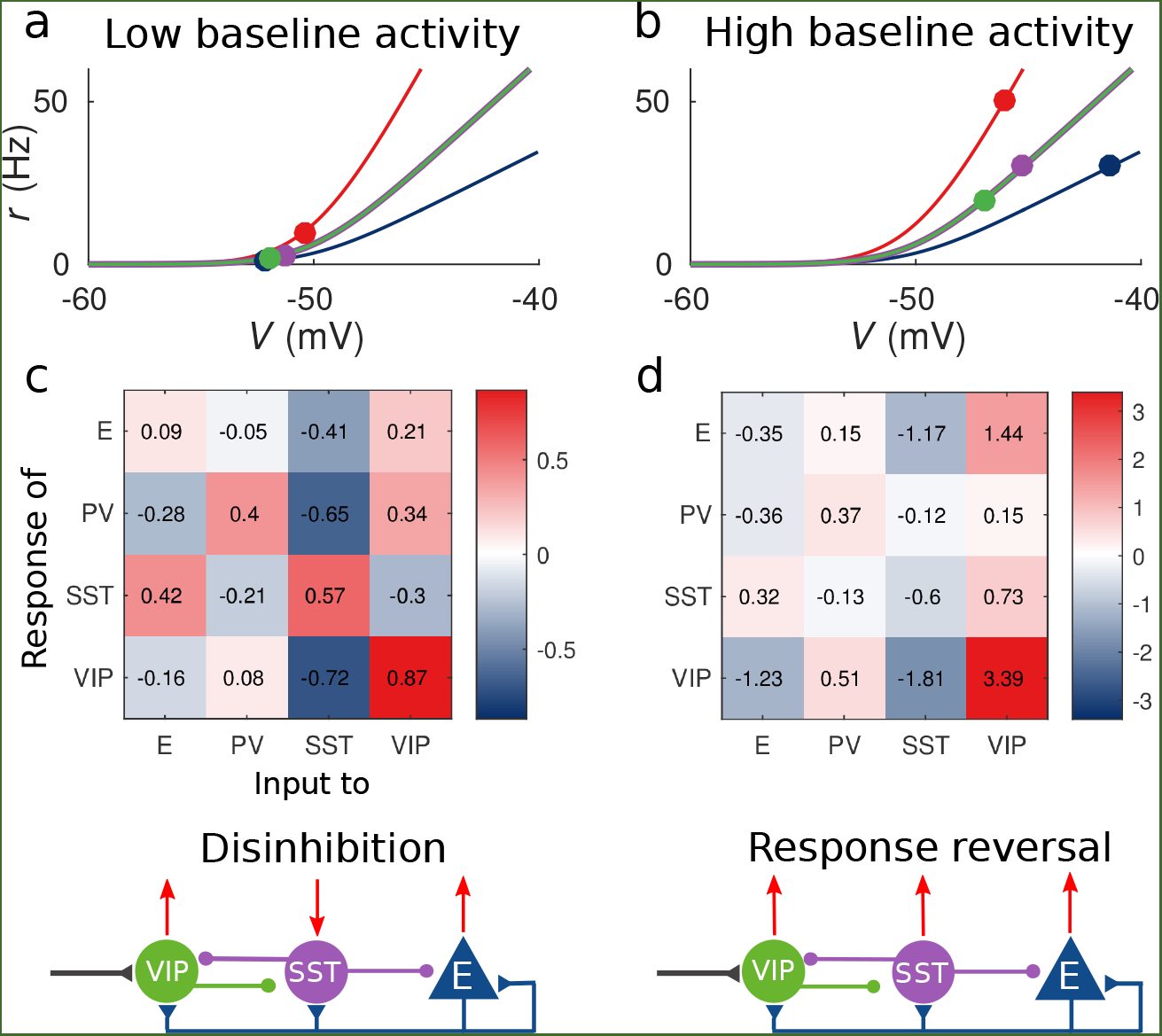
Response matrix and disinhibition vs. response reversal regime. (a-b) Tuning curves for the different populations and baseline activity in both scenarios (low and high). In the low baseline activity scenario (a) all populations are below threshold (flat part of the fI curve), instead in the high baseline activity scenario (b) all populations are above threshold, where small changes in input result in large changes in rate. (c-d) Response matrices for the two scenarios. In (c) the response of SST to external excitation of VIP is negative, while the responses of E and PV are positive. This corresponds to the disinhibition regime. In (d) the responses of all populations to external excitation of VIP are positive, in particular, the response of SST is reversed with respect to (c) corresponding to the response reversal regime.

**Figure 3:**
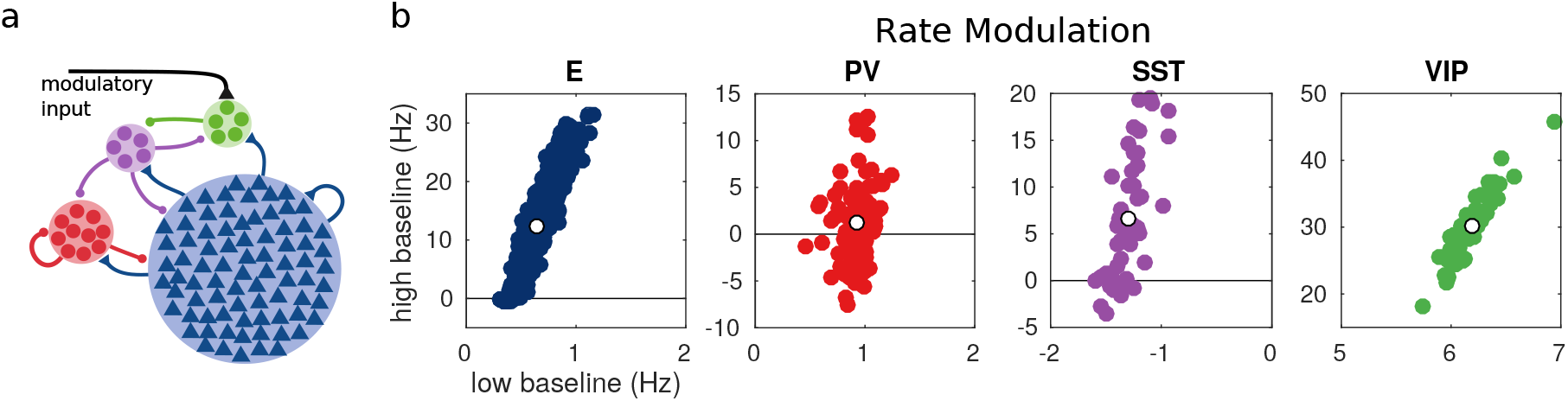
Random network model. (a) Schematic of the model. Each population is compsed of several rate units and the connectivity between units is random with probabilities extracted from experimental data in the literature. (b) Rate modulation (rate after the onset of the modulatory current minus baseline rate) for low and high baseline activities. Each colored point corresponds to one unit. Unit responses are very variable and, in particular within the same population different units might have responses with different sign. White points correspond to the population average. Despite the variability of individual responses the population average corresponds to the population responses in the single unit model in figure 1.

### Random network model

Experimental recordings showed a great diversity across neural responses even when recording from the same type of cells. Although this diversity can have many origins, such as different cell subtypes, we proposed that random connectivity alone is sufficient to explain it. To do so we develop an extension of our model where each population is composed of multiple rate units and where the probability that one connection exists from one unit to another depends on the populations of the presynaptic and postsynaptic units according to data extracted from [9, 25] (see methods for details).

For each unit we measure the rate modulation (rate during top-down modulation minus baseline activity) for the different baselines. If the rate modulation is positive it means that the neuron is more active in the presence of the modulatory current and vice versa. In 3b we show scatter plots of the rate modulation in under the low baseline condition versus the rate modulation under the high baseline condition for each unit. These simulations reveal that due to the heterogeneity in the connectivity, the behavior of individual neurons can be quite variable while the population average still corresponds to the behavior of the population based model. This variability can result in cells within the same population having responses with opposite sign, as has been observed to be the case in mouse V1 [3, 24, 27] and A1 [12]. In addition variability might also have further implications for gating of signals, since variability in inhibitory cells has been proposed to modulate the response gain of neural circuits [19].

### Simulation of V1 accounts for experimental measurements

Our framework allows us to easily understand the counterintuitive behavior of V1 during locomotion [3, 4, 24]. Different levels of visual stimulation result in different baseline activities and in this case top-down modulation is triggered by locomotion.

To model visual input we use external currents. In the case of size-varying gratings this input has two sources: thalamic input that targets excitatory cells and cortical input that targets SST cells. In order to reproduce the surround suppression effect [2, 23] excitatory cells have a small receptive field and therefore receive center input and SST cells have a large receptive field and receive surround input (see methods for details).

Figure 4b shows the response reversal phenomenon when a weak visual stimulus is presented. Before the visual stimulation the SST has higher activity for immobility than for locomotion, by contrast, when the visual stimulus is presented, the activity of the SST population is higher for locomotion. In figure 4c we show the experimental data from [24] for three different experimental conditions (darkness, gray screen and grating) and in figure 4d our simulations of V1 under the same conditions. Similarly figure 4e shows the experimental data from [3] for gratings of different sizes and 4f shows the behavior of our model.

**Figure 4:**
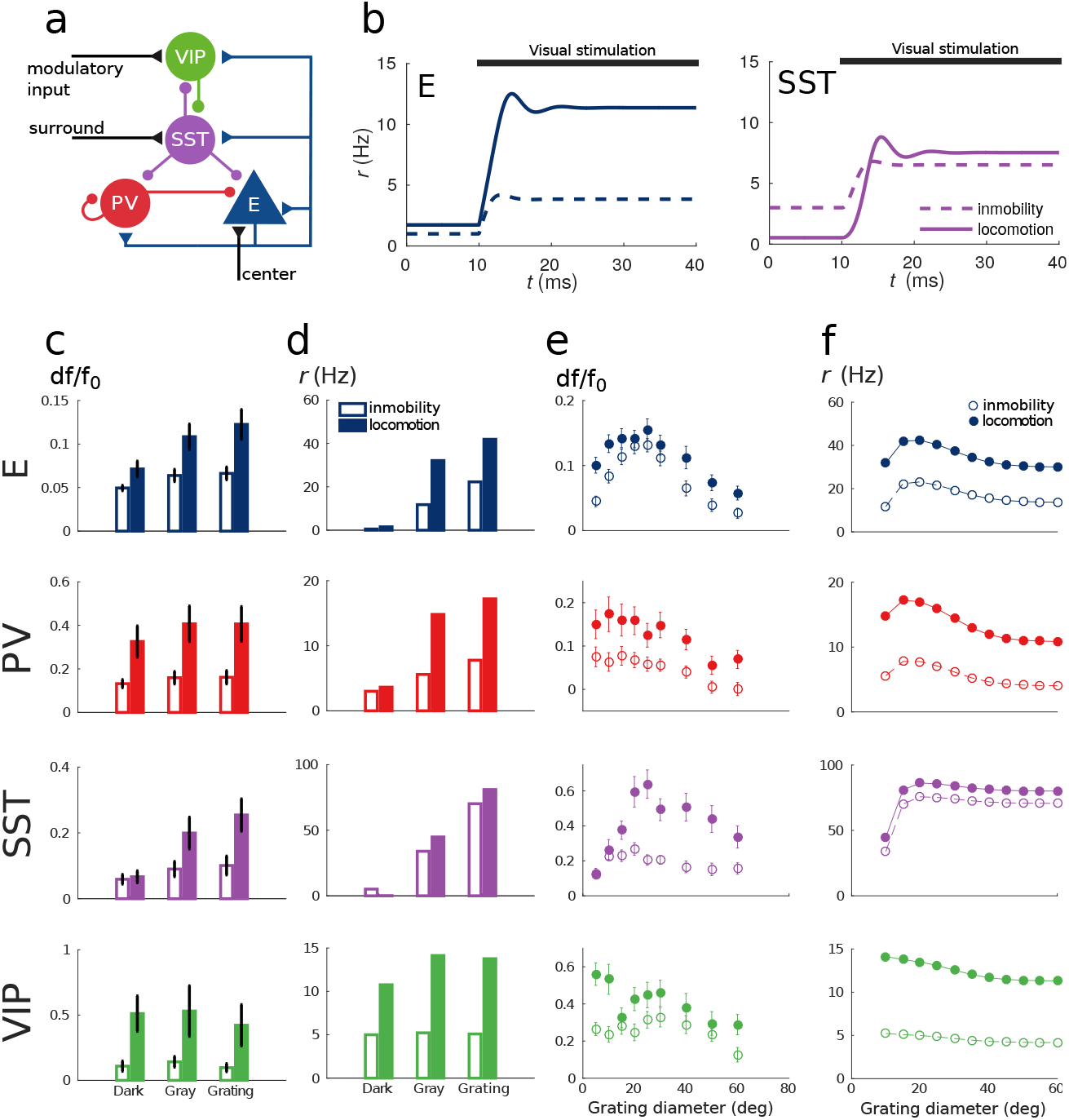
Model of mouse V1 behavior. (a) Schematic of the microcircuit. Visual input targets E and SST cells. Behavior related top-down modulation targets VIP cells. (b) Response of E and SST populations when a weak visual stimulus (6 deg) is presented for locomotion and immobility. The E population always shows a higher response with locomotion. On the other hand, before the visual stimulation the SST population has higher activity for immobility than for locomotion and when the visual stimulus is presented, the activity of the SST population is higher for locomotion. (c) Relative change in calcium fluorescence for three levels of visual stimulation (darkness, gray screen and grating) and two behavioral states: immobility (empty bars) and locomotion (filled bars) extracted from [Pakan et al. 2016]. (d) Rates (in Hz) of the populations in the V1 simulation for the same conditions as in (c). (e) Relative change in calcium fluorescence for gratings of diameters ranging from 10 deg to 60 deg for the two behavioral states: immobility (empty dots) and locomotion (filled dots) extracted from [Dipoppa et al. 2016]. (f) Rates (in Hz) of the populations in the V1 simulation for the same conditions as in (e). Comparison of (c) with (d) and (e) with (f) shows that our simulations reproduce qualitatively the activity of neural populations in mice V1. Namely the activity of all populations is higher during locomotion than during immobility whenever there is visual stimulation and for E, PV and VIP also in the absence of visual stimulation. Our model shows a decrease in activity of SST during locomotion as reported in the experiments (the change in activity of the SST population in darkness in (b) is not statistically significant). Our model also exhibits surround suppression for all populations. The quantitative differences might be related to the fact that changes in calcium fluorescence are not proportional to changes in rate.

Our simulations of this V1 circuit model reproduce the phenomena described in the literature: in darkness, the activities of excitatory, PV and VIP populations increase during locomotion whereas the activity of the SST population decreases with respect to the activity during immobility [3, 4]. In the presence of visual stimulation the activities of all populations, including SST, increase during locomotion [3, 24].

To show that our results do not rely on a fine tuning of the connectivity parameters or even on certain details of the microcircuit structure we have run the model with several connectivity matrices and perturbations of them (figure S1). We have also considered other microcircuit structures to account for the differences between studies ( [25] reports projections from PV to VIP and [9] from PV to SST) and we also consider thalamic input to PV (figure S2). In all these cases, the results were consistent with our original findings.

## Discussion

We developed a model that reproduces two counterintuitive phenomena observed in mouse cortex. First, in certain cases the activation of VIP cells results in an overall positive response of the SST population [3, 24]. Second, the sign of the SST population response to excitation of VIP cells depends on the baseline activity of the circuit [3, 4]. Two features of the system lead to this behavior: the presence of multiple interneuron populations and the nonlinearity of f-I curves.

We explained heuristically the response reversal by closely looking at transient dynamics of the circuit. One experimentally-testable prediction of our analysis is that in the response reversal regime, the overall SST population response to top-down modulation should initially decrease and later increase until reaching a higher rate than the baseline.

Based on our model we introduced the response matrix *M*, which is a comprehensive framework to understand counterintuitive steady state responses. It provides explicit information about the contribution of each individual connection. For example by looking at the elements in *M_SV_* (see table 3), one can readily see that if the recurrent excitation between pyramidal cells is not large enough, *M_SV_* can only be negative and therefore response reversal of SST would not happen. Another example is that if both SST and VIP populations have high baseline activities and if the SST-VIP-SST loop is strong enough, *M_EE_* can be negative, i.e. the excitatory population can have a negative response to excitatory input (see table 3 for the explicit expression of *M_EE_*). If the connections between the SST and the VIP populations are removed (or weakened) or if their baseline activities are sufficiently lowered *M_EE_* will always be positive. This constitutes another interesting prediction that can be experimentally tested.

Our calculations also revealed sign correlations between entries of *M*, for example *M_SV_* and *M_SS_* have opposite signs for any connectivity matrix (given the microcircuit) and for any baseline activity. This predicts that in the regime where SST activity has a positive response to excitatory input targeting VIP, SST has to have a negative response to external input targeting SST. This prediction means that increased grating size, which provides extra excitation to the SST population [2], should actually decrease the SST activity, as observed in both data [3] and our model but not in previous experiments [2].

The analysis of the response matrix shows that for the given microcircuit structure all terms of the matrix can be positive or negative. This is not the case for a network with one excitatory (E) population and only one inhibitory (I) population [23, 30]. In that case *M_EE_* and *M_IE_* are always positive, *MEI* is always negative and only *M_II_* can have both signs. In this sense, having more than one inhibitory population results in a much more versatile network.

Our approach constitutes a general conceptual framework in which previous work can be better understood [18, 23, 30]. It provides a parsimonious yet powerful explanation to surprising observations of interneuronal circuits in V1 [3, 13, 24] without assuming top-down excitatory inputs targeting SST or PV neurons. Furthermore it could be extended to explain similar phenomena observed in A1 [12, 29]. In addition it is in line with experimental results that show that VIP interneurons play an important role in cortical activity modulation [7, 8, 20].

We have shown that similarly to the now well-known paradoxical effect that the presence of a single inhibitory neuron type can cause [23, 30], the presence of multiple types of interneurons has an even stronger impact on the activity of neural circuits. We have also exposed the effect of nonlinearity of the f-I curve. Our analysis suggests that in a circuit with multiple populations, the most interesting circuit behavior is found when spontaneous baseline activity is close to threshold since in that regime responses will change the most with small changes in population rates. These two features signifficantly broaden the richness of the dynamics of cortical circuits and enhance their usefulness for cognitive and behavioral computations. We conclude that computational models and mathematical analysis are critical to fully understand the dynamics of neural circuits underlying behavior, especially when several types of interneurons are involved as intuition alone may be misleading and provide erroneous predictions on such circuits.

## Methods

### Firing rate based population model

The state of the system is characterized by the rates *r_i_*. To model the average rate of each population we use a function of the input *V_i_* as the one introduced in [1]

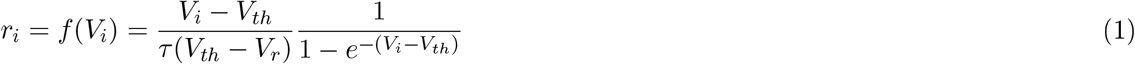

where *V_th_* = −50 mV and *V_r_* = −60 mV are the threshold and reset potentials respectively and τ is the membrane time constant. *V_i_* is the average input to each of the populations and is given by

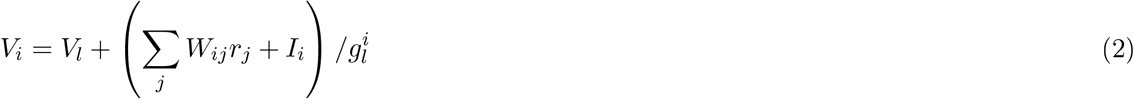

where *V_l_* = −70 mV is the reversal potential and *g_l_* is the membrane conductance. *W* is the connectivity matrix and therefore 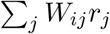 is the recurrent local input. *I_i_* is the external input current. The rate dynamics are given by

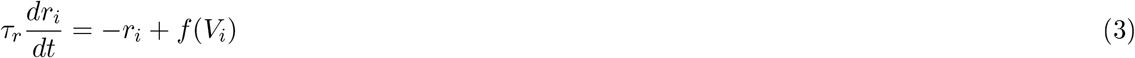

where *τ_r_* = 2 ms [5]. Since the parameters of the f-I curve are population dependent (see table 2), different populations will have different rates for the same input. The nonlinearity of the f-I curve has very important consequences. Namely, for low input *f*(*V_i_*) is almost flat, and therefore changes in the input will have almost no effect on the rate. By contrast, for strong input *f*(*V_i_*) tends asymptotically to a straight line with slop 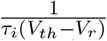 and changes in the input will elicit a large change in the rate. As we will show later, this feature is key to reproduce the response reversal observed in the experiments.

The connectivity matrix *W* is generated by rejection sampling, i.e. by generating random matrices that have the microcircuit structure (inhibitory and excitatory connections) and selecting the ones that produce the desired responses. The simulations of figures 1 and 2 where done with the connectivity matrix given in table 1.

**Table 1:**
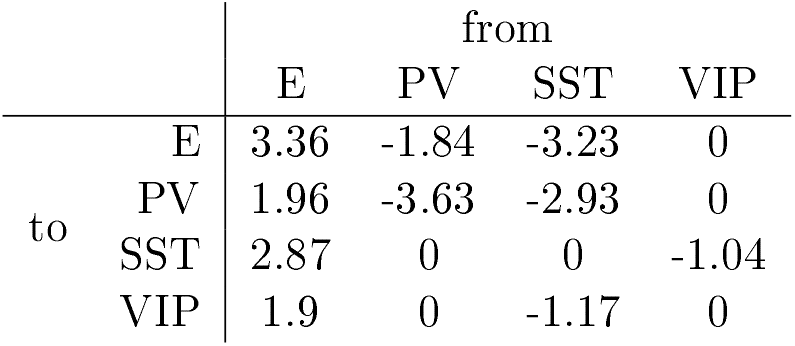
Connectivity matrix.

|  |  | from |  |  |  |
| --- | --- | --- | --- | --- | --- |
|  |  | E | PV | SST | VIP |
| to | E | 3.36 | -1.84 | -3.23 | 0 |
|  | PV | 1.96 | -3.63 | -2.93 | 0 |
|  | SST | 2.87 | 0 | 0 | -1.04 |
|  | VIP | 1.9 | 0 | -1.17 | 0 |

Behavioral state is modelled with a constant top-down modulatory current of 10 pA that targets VIP cells. We also include a constant background input so that in the absence of the top-down modulatory current, the E, PV, SST and VIP populations will have spontaneous average rates of 1, 10, 3 and 2 Hz respectively for the low baseline activity scenario and 30, 50, 30 and 20 Hz for the high baseline activity.

### Response matrix and response reversal

In order to characterize the response of a population to external excitatory input to the network we calculate how its rate will change for a small change in external input. We focus on stationary states *r_i_* = *f*(*V_i_*). If we apply a small perturbation to the external input *δI_i_*, the network will reach a new stationary state

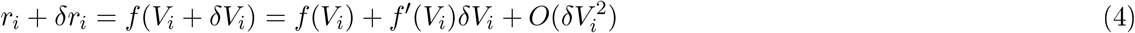

where *f*′(*V_i_*) is the derivative of *f* with respect to *V* and

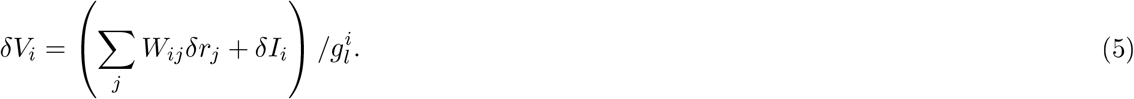

Since *r_i_* = *f*(*V_i_*), when linearize *f* around *V* and ignore terms of order *δV*^2^ and higher we obtain the following self-consistent equation

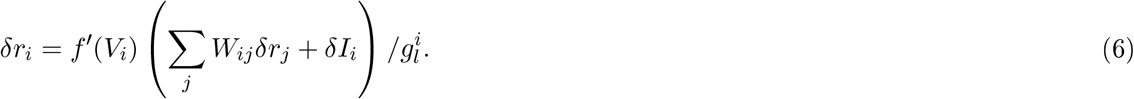

We define the entries of response matrix as the derivative 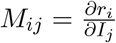, which can be obtained from the limit *δI_j_* → 0 in the system of equations given by (6) and in matrix form can be written as

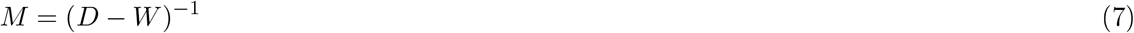

where *D* is a diagonal matrix with entries *D_ii_* = *g_l,i_*/*f*′(*V_i_*). As it was explained in the results section, the nonlinear behavior of the terms *D_ii_* is essential to explain the response reversal regime. *D_ii_* becomes arbitrarily large as *V_i_* → -∞ and decreases monotonically to 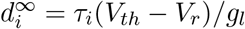 when *V_i_* → ∞.

In table 3 we give the explicit formulas to all the entries of the response matrix in terms of the entries of the connectivity matrix *W* and *D* (we denote *w* = |*W*|, *d_i_* = *D_ii_* and *C* = det(*D* − *W*) ^−1^). Note that, because of the complex interactions in the network, the sign of *M_ij_* is never determined exclusively by that of *W_ij_*.

### Random network model

We consider a network with 800 E units, 100 PV units, 50 SST units and 50 VIP units. Each unit within a population has the same f-I curve with the parameters in table 2. The probabilities *p_ij_* of a connection from each unit in population *j* to each unit in population *i* are estimated from data [9, 25] adnare given in table 4.

**Table 2:**
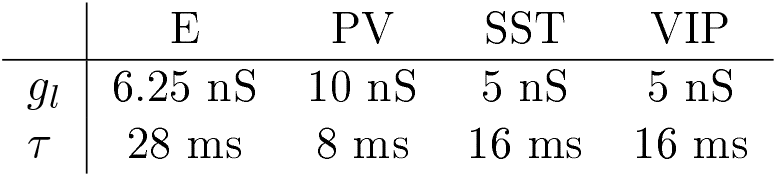
Population dependent parameters.

|  | E | PV | SST | VIP |
| --- | --- | --- | --- | --- |
| $g_l$ | 6.25 nS | 10 nS | 5 nS | 5 nS |
| $\tau$ | 28 ms | 8 ms | 16 ms | 16 ms |

**Table 3:** Entries of the respone matrix.

|  |
| --- |
| $M_{EE} = C(w_{PP} + d_P)(d_S d_V - w_{SV} w_{VS})$ |
| $M_{PE} = C(w_{PE}(d_S d_V - w_{SV} w_{VS}) - w_{PS}(w_{SE} d_V - w_{SV} w_{VE}))$ |
| $M_{SE} = C(w_{PP} + d_P)(w_{SE} d_V - w_{SV} w_{VE})$ |
| $M_{VE} = C(w_{PP} + d_P)(w_{VE} d_S - w_{SE} w_{VS})$ |
| $M_{EP} = -C w_{EP}(d_S d_V - w_{SV} w_{VS})$ |
| $M_{PP} = -C((w_{EE} - d_E)(d_S d_V - w_{SV} w_{VS}) + w_{ES}(w_{SE} d_V - w_{SV} w_{VE}))$ |
| $M_{SP} = -C w_{EP}(w_{SE} d_V - w_{SV} w_{VE})$ |
| $M_{VP} = -C w_{EP}(w_{VE} d_S - w_{SE} w_{VS})$ |
| $M_{ES} = -C d_V(w_{ES}(w_{PP} + d_P) - w_{EP} w_{PS})$ |
| $M_{PS} = -C d_V(w_{ES} w_{PE} - (w_{EE} - d_E) w_{PS})$ |
| $M_{SS} = -C d_V((w_{EE} - d_E)(w_{PP} + d_P) - w_{EP} w_{PE})$ |
| $M_{VS} = -C(w_{VE}(w_{ES}(w_{PP} + d_P) - w_{EP} w_{PS}) + w_{VS}((w_{EE} - d_E)(w_{PP} + d_P) - w_{EP} w_{PE}))$ |
| $M_{EV} = C w_{SV}(w_{ES}(w_{PP} + d_P) - w_{EP} w_{PS})$ |
| $M_{PV} = C w_{SV}(w_{ES} w_{PE} - (w_{EE} - d_E) w_{PS})$ |
| $M_{SV} = C w_{SV}((w_{EE} - d_E)(w_{PP} + d_P) - w_{EP} w_{PE})$ |
| $M_{VV} = C(w_{ES}(w_{ES}(w_{PP} + d_P) - w_{EP} w_{PS}) - d_S((w_{EE} - d_E)(w_{PP} + d_P) - w_{EP} w_{PE}))$ |

**Table 4:** Connection probabiliries for the random network model.

|  |  | from |  |  |  |
| --- | --- | --- | --- | --- | --- |
|  |  | E | PV | SST | VIP |
| to | E | 0.02 | 1 | 1 | 0 |
|  | PV | 0.01 | 1 | 0.85 | 0 |
|  | SST | 0.01 | 0 | 0 | -0.55 |
|  | VIP | 0.01 | 0 | 0.5 | 0 |

The strengths of the connections are rescaled so that the average input of a unit in population *j* from all units in population *i* is *Wij*. Top-down modulatory current and background input is identical to all units within the same population.

### Mouse V1 model

In the simulations of V1 activity we use the connectivity matrix given in table 5.

**Table 5:** Connectivity matrix for the mouse V1 model.

|  |  | from |  |  |  |
| --- | --- | --- | --- | --- | --- |
|  |  | E | PV | SST | VIP |
| to | E | 3.30 | -3.48 | -2.98 | 0 |
|  | PV | 1.73 | -4.25 | -1.07 | 0 |
|  | SST | 3.50 | 0 | 0 | -4.51 |
|  | VIP | 0.53 | 0 | -0.13 | 0 |

We model thalamic input with an external excitatory current that targets E and SST cells. In the experiments in [3, 24] the authors consider three levels of visual stimulation which are: darkness, gray screen and grating. To model darkness condition we assume a total absence of visual stimulation (therefore *I_E_* = 0 pA, *I_S_* = 0 pA). For gray screen we use a small input current to the excitatory population (*I_E_* = 50 pA, *I_S_* = 0 pA). Finally to model different grating diameters the value of the input is a sigmoid function of the grating diameter *θ*:

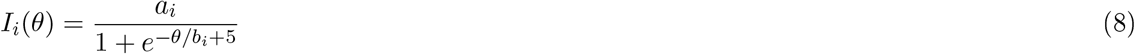

where *b_E_* = 2, *b_S_* = 6, *a_E_* = 100 pA, *a_S_* = 20 pA. With this parameters E cells receive center input (input saturates for diameters ~20 deg) and SST cells receive surround input (input to SST saturates for diameters of ~60 deg) [3].

To demonstrate that our results do hold for a wide range of connectivity matrix and do not have to be ne tuned, we simulate several different connectivity matrices that produce the same qualitative behavior. We also make perturbations of these matrices by multiplying each entry by a random variable uniformly distributed in the interval [0.9; 1.1]. This amounts to randomly modifying each connection within ±10% of its original value (see figure S1).

In the alternative models of figure S2 where visual stimulus input also targets PV cells, we use *I_P_* = 0 pA for darkness, *I_P_* = 10 pA for gray screen and *b_P_* = 2, *a_P_* = 20 pA for gratings.

---

**Figure S1:**
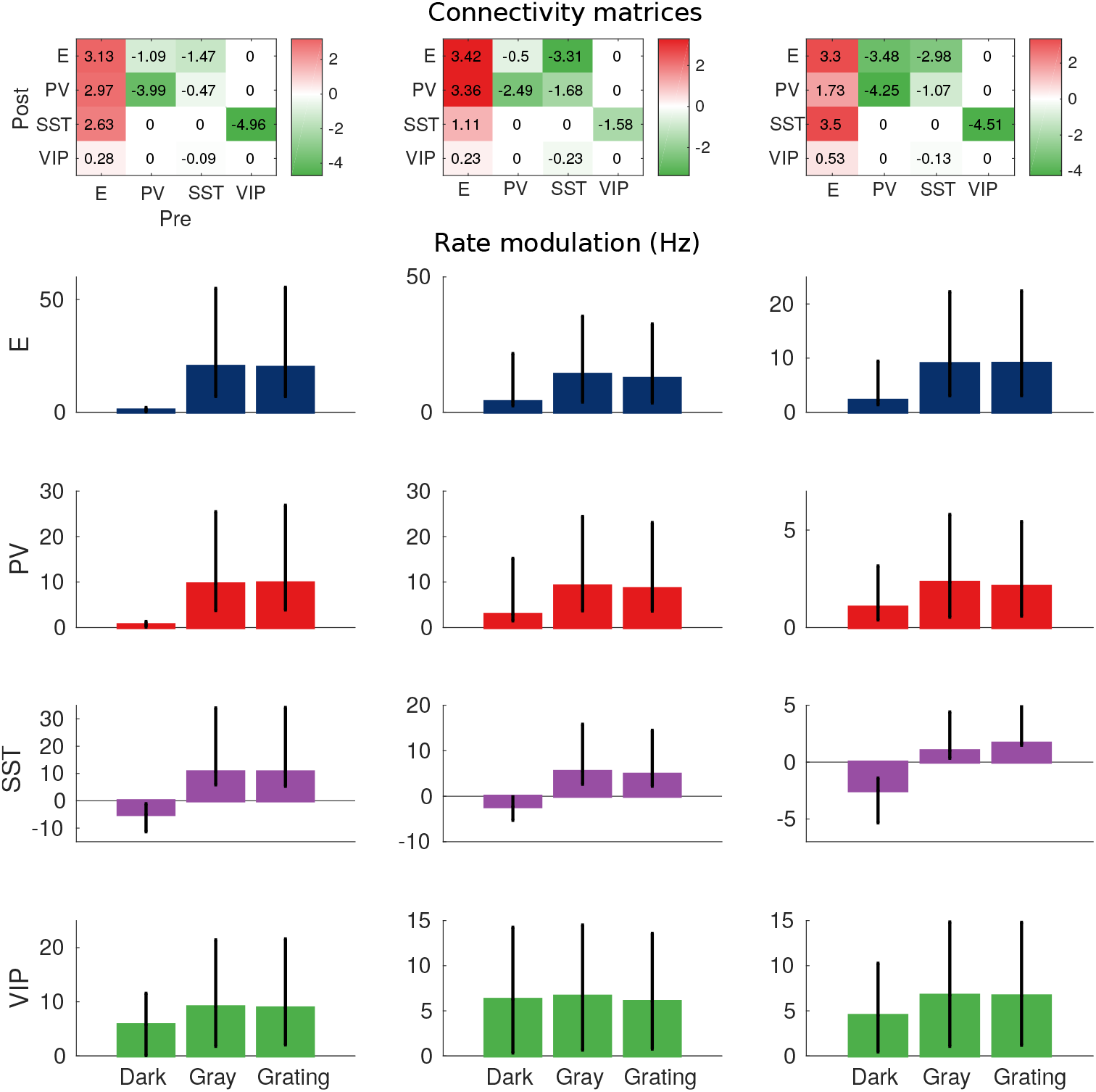
Robustness of the behavior. Top: Example of three connectivity matrices that have the same qualitative behavior. Bottom: rate modulation (rate during locomotion minus rate for immobility). Each bar corresponds to the average rate modulation of 20 random perturbations of the matrices on the top where each entry has been multiplied by a random variable uniformly distributed in [0.9; 1.1], which corresponds to random changes of up to ±10%. Error bars correspond to the minimum and maximum rate modulations of the 20 realizations. Despite quantitative variations, the qualitative behavior is always the same: rate modulation of SST population in darkness is always negative; rate modulation for all other cases is always positive.

**Figure S2:**
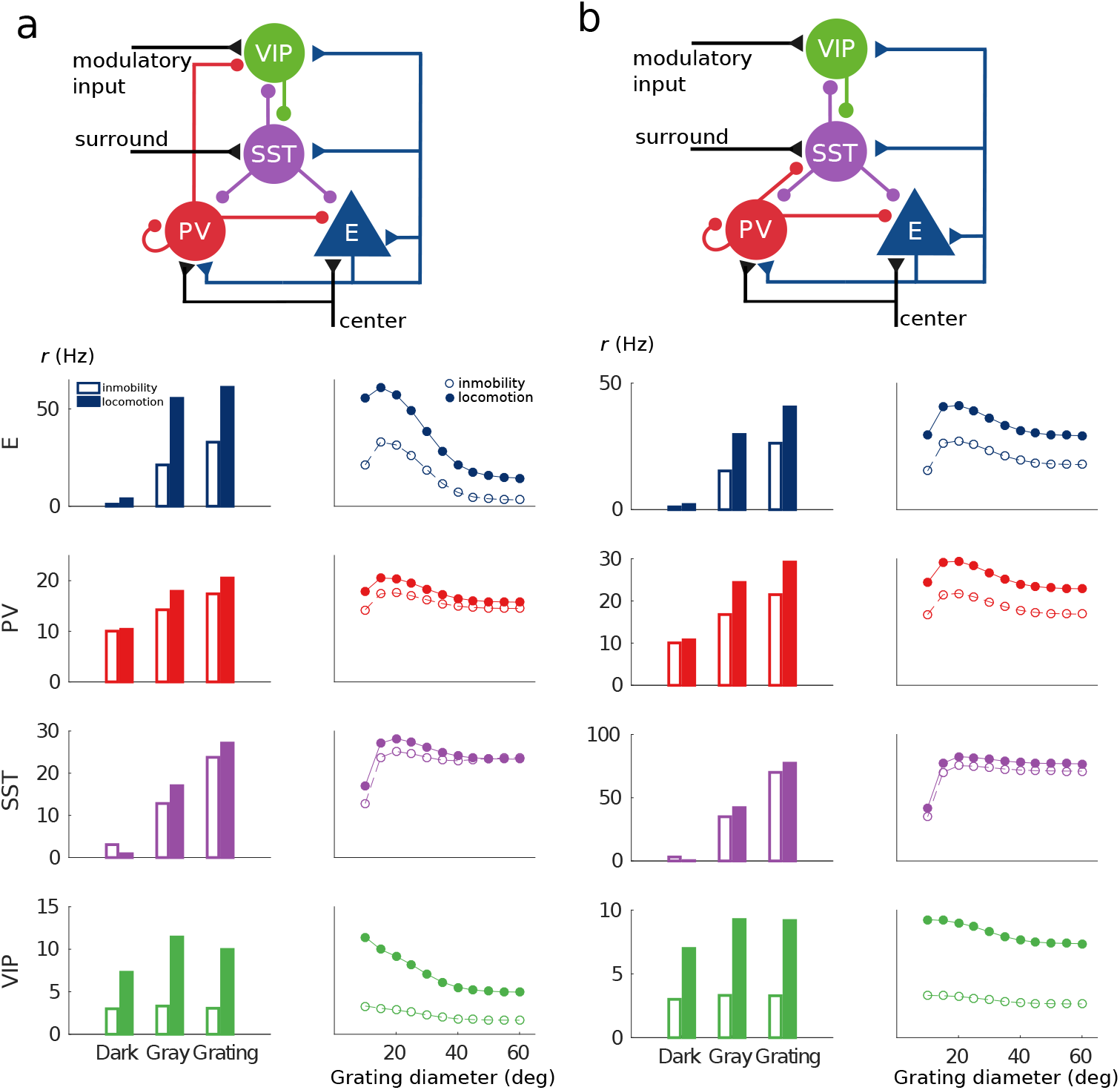
Two alternative microcircuits with visual input targeting E, SST and PV populations and PV to VIP (a) and PV to SST (b) connections.

